# Comparative transcriptome analysis reveals higher expression of stress and defense responsive genes in dwarf soybeans obtained from the crossing of *G. max* and *G. soja*

**DOI:** 10.1101/455923

**Authors:** Yong-Wook Ban, Neha Samir Roy, Heejung Yang, Hong-Kyu Choi, Jin-Hyun Kim, Prakash babu, Keon-Soo Ha, Jin-Kwan Ham, Kyong Cheul Park, Ik-Young Choi

## Abstract

Plant height is an important component of plant architecture and significantly affects crop breeding practices and yield. We obtained a few segregated dwarf soybeans in the populations derived from the crossing of *Glycine max* var. Peking and *Glycine soja* var. IT182936 in an F5 RIL population. These dwarf soybeans may be useful genetic resources for plant breeders, geneticists and biologists. We attempted to find differentially expressed genes to classify and understand the regulation of genes related to plant growth in mutant dwarf soybeans, which appeared in the F5 generation. Using the Illumina high-throughput platform, transcriptomes were generated and compared among normal and dwarf soybeans in triplicate. We found complex relationship of the expressed genes to plant growth. There are highly significantly up-/downregulated genes according to the comparison of gene expression in normal and dwarf soybeans. The genes related to disease and stress responses were found to be upregulated in dwarf soybeans. Such over-expression of disease resistance and other immune response genes was targeted to understand how the immune genes regulate the response of plant growth. In addition, photosynthesis-related genes showed very low expression in dwarf lines. The transcriptome expression and genes classified as related to plant growth may be useful resources to researchers studying plant growth.

## Introduction

Dwarfism is a desirable trait in crop plants, as it prevents lodging and, in most cases, makes the plant study [1]. Dwarfism was one of the major contributors to the ‘green revolution’, which brought about a boom in agricultural production [2]. However, plant dwarfism cannot always be linked to increased yield. Plants sacrifice their growth for different defense responses [3]. Several factors may affect plant height such as light and soil nutrients [4], drought [5, 6], waterlogging [7], floods [8], cold temperatures [9], infection [10, 11], and herbivory [12]. In soybeans, some high-yielding and lodging-resistant dwarf cultivars have been developed [13-16] ; however, there has not yet been any comparative transcriptome study dissecting the genetic factors involved in dwarfism in soybean recombinant inbred lines (RILs).

Dwarfism in plants is basically due to the cessation of cell elongation [17] and inhibited cell division [18] brought about by one or more of the aforementioned factors. Optimum levels of plant growth hormones (PGRs), particularly, gibberellin and auxin, and their interactions are known to play important roles in plant growth [19, 20]. Interestingly, plants under defense pressure are known to redirect their resources to produce one or more of the defense response metabolites (salicylic acid (SA), jasmonates (JA) etc.), thereby reducing overall growth [3]. A transcriptome analysis reported that multiple genes involved in hormone biosynthetic pathways were influenced in dwarf soybean mutants [21]. In another study, knockout of a gene of the peroxidase superfamily reportedly brought about dwarfism in soybean plants [22].

In the present study, we performed a comparative transcriptome analysis of three normal and three dwarf soybean F5 RILs derived from the cross between *G. max* and *G. soja*. Numerous differentially expressed genes (DEGs) were identified using different bioinformatic tools, and putative associations between major DEGs were established and discussed.

## Materials and Methods

### Plant materials

The soybean hybrid developed from *Glycine max* var. Peking and *G. soja* var. IT182936 was used for the development of F5 RILs. The soybeans used in this experiment were harvested in Chuncheon city (Gangwon-do, South Korea). F5 RILs with distinctly different phenotypes (based on height) were used, which included tall or normal phenotypes: 1214-1 (N1), 1273 (N2) and 1321 (N3); and dwarf: 1214-2 (D1), 1265-2 (D2), and 1265-3 (D3) (Fig 1). The leaves were collected from each sample before flowering, frozen immediately in liquid nitrogen and stored at -80°C.

**Fig 1.**
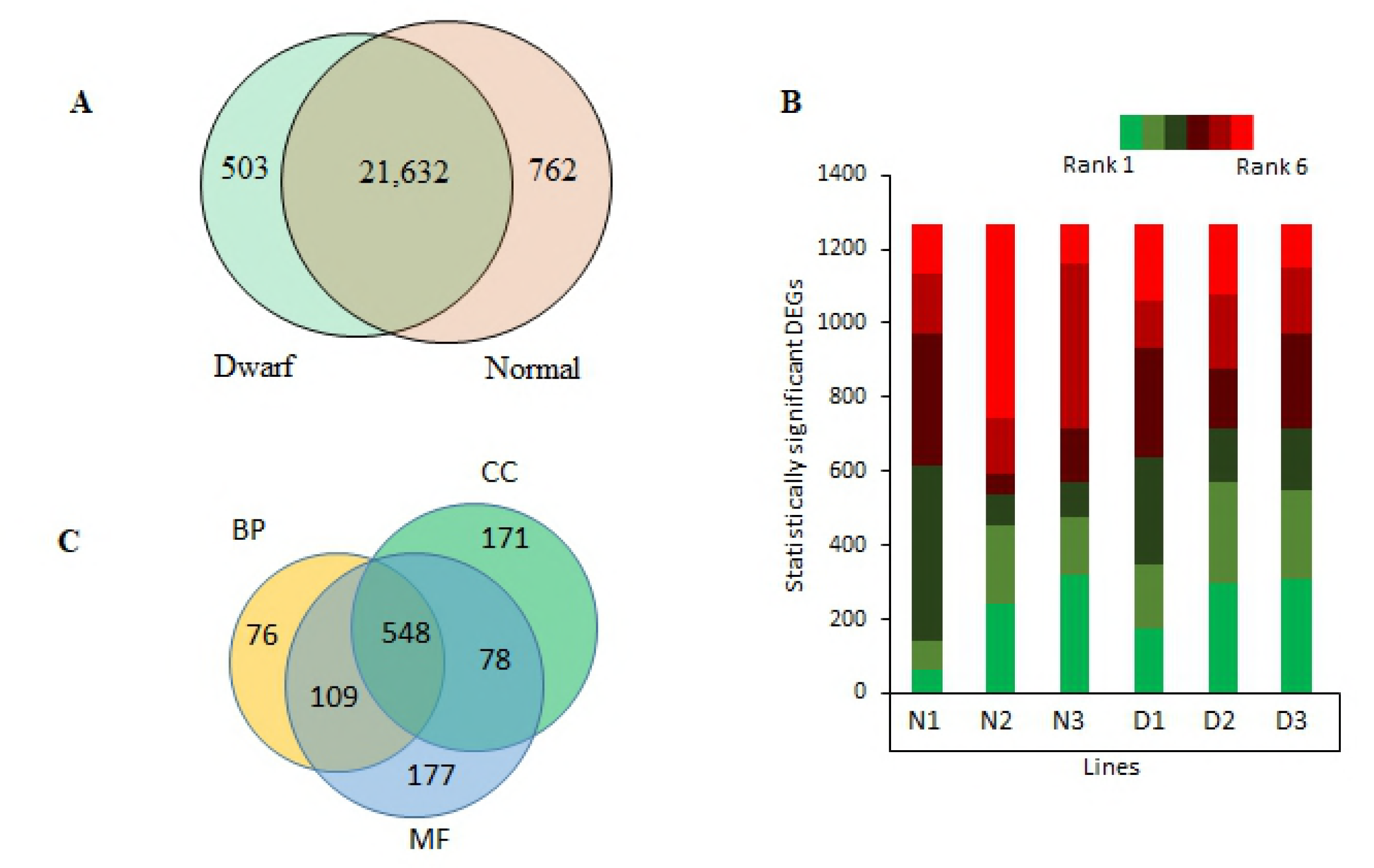
RIL lines used in the current study

### RNA isolation and cDNA synthesis for RNA sequencing

The total RNA was purified from leaves of each of the six RIL lines (N1-3, D1-3) three weeks after germination using the RiboPure kit (Applied Biosynthesis, Foster City, CA, USA) following the manufacturer’s protocol. DNA digestion was performed with DNase **I** (Sigma, St. Louis, MO, USA), and the total RNA was quantified by absorption of light at A260 and the quality checked by analyzing the ratios A230/260 and A260/280 using a NanoDrop^®^ 2000 Spectrophotometer (Thermo Scientific, Wilmington, DE, USA). Paired-end sequencing was performed with a HiSeq 2500 (Illumina, San Diego, CA, USA) at the National Instrumentation Center for Environmental Management (NICEM, Seoul National University, Seoul, South Korea). The raw reads were cleaned by filtering out adaptor-only nucleotides and trimming of adaptor nucleotides, empty nucleotides (N in the end of reads), short reads (<59 bp) and low quality nucleotides (reads containing more than 50% bases with Q-value ≤20) using trimmomatic [23]

### DEG (differentially expressed genes) analysis

The Williams 82 Assembly 2 Annotation 1 (Wm82.a2.v1) coding sequence provided on the SoyBase website (https://soybase.org/GlycineBlastPages/blast_descriptions.php) was used as the reference gene set. The expression value for each gene was calculated as FPKM (fragments per kilobase of transcript per million mapped reads). Hisat2 [24] was used for the mapping, and the number of mapped reads for each unigene set was obtained from the mapping result file for each sample using Samtools [25]. The log2-fold change value was calculated as the normalized FPKM value of the expression level of the sample to be compared for analysis. The up- and down-regulation analyses were performed using Fischer’s exact test (p ≤0.05). Additionally, the false discovery rate (FDR) was calculated for the significantly differentially expressed genes (DEGs). When the DEGs showed relatively higher FPKM values in tall RILs, they were regarded as upregulated but when this case occurred in dwarf RILs, the DEGs were regarded as downregulated.

### HPLC analytical conditions

For HPLC, an Agilent system (Agilent, Santa, CA, USA) comprised of a pump (Agilent 1260 Quat Pump VL DEAB818744), an auto sampler (Agilent 1260 ALS DEAAC38064), a column oven (Agilent 1260 TCC DEACN40488), and UV detector [Agilent 1260 MWD VL DEAAZ01707] were used. HPLC analysis was conducted on a HECTOR C18-M column, C18 – M51002546 (4.6 mm I.D. × 250 mm, 4.5-μm pore size). The mobile phase consisted of 0.1% FA water (A) and methanol (B), operating at a flow rate of 1.0 mL/min. The HPLC linear gradient profile was as follows: 40% B at 0 min, 40% B at 0 – 25 min, 40 - 100% B at 25 – 25.1 min, 100% B at 25.1 – 35 min, 100 – 40% B at 35 – 35.1 min, and 40% B at 35.1 – 40 min. The injection volume was 10 μL. The column temperature was set to 30°C. For the determination of each standard compound, UV spectra were evaluated at 300 nm. Salicylic acid was purchased from Sigma-Aldrich (Korea). The purities of the standard were above 98%. HPLC-grade methanol and water were purchased from TEDIA (Fairfield, OH, USA). Formic acid (FA) was purchased from DAE JUNG (Korea). Standard stock solutions of salicylic acid (500 μg/mL) were prepared in methanol. These standard stock solutions were diluted to 5 different concentrations by methanol to establish the calibration curves. The 100% metabolic extracts from each of three dwarf and normal F5 RILs and the initial parents were weighed and dissolved in methanol at the rate of 20.00 mg/mL. Before HPLC analysis, all of the sample solutions were filtered through a 0.45-μm membrane independently.

## Results

### RNA-seq analysis of soybean RILs

Over the generations, RIL lines of *G. max* and *G. soja* had tall plants, in most cases, that further segregated to tall and dwarf phenotypes, while dwarf plants consistently produced progenies of their own phenotype (data not shown), giving an initial clue that the dwarf phenotype involves recessive gene/s. The tall plants had longer and thinner internodes, broader leaves and viney growth resembling *G. soja*, while dwarf plants had shorter and thicker internodes, narrow leaves, and *G. max*-like upright growth (Fig 2). Other than these traits, almost all morphological characteristics, such as color of seeds and flowers, in the dwarf lines were similar with other lines. To gain deeper insight into dwarfism, a comparative transcriptome analysis was performed between normal and dwarf plants, and significantly up- and downregulated genes were screened. In total, >105 million (M) RNA-Seq reads were generated from the tall and dwarf RILs. After trimming, a total of 65 M high quality reads were obtained (Table 1).

**Fig 2.**
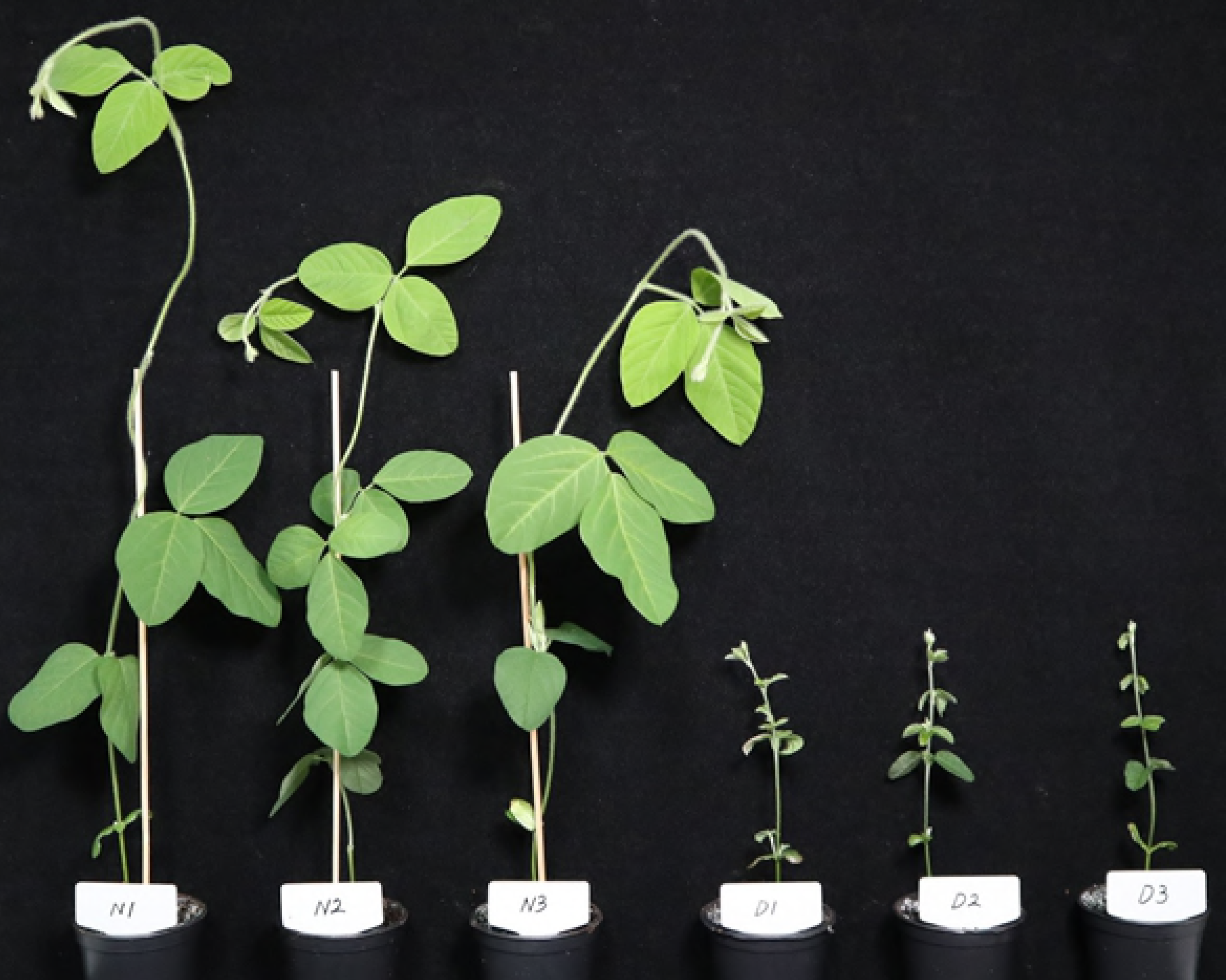
Growth behavior and general morphology of the normal and dwarf F5 RILs (4 weeks after sowing). The dwarf types displays much shorter plant height than other types.

**Table 1.** Summary of total RNA-seq data from soybean F5 RILs.

| Sample | Reads | Length | Q30(%) | Q20(%) | GC Ratio(%) |
| --- | --- | --- | --- | --- | --- |
| N<sub>1</sub> | 22,113,816<br>(12,244,292) | 3,339,186,216<br>(1,790,053,878) | 92.34 (98.03) | 95.18 (99.22) | 49.91 (48.59) |
| N<sub>2</sub> | 12,898,488<br>(7,664,212) | 1,947,671,688<br>(1,139,481,523) | 97.15 (98.99) | 98.38 (99.64) | 51.28 (49.94) |
| N<sub>3</sub> | 20,513,596<br>(9,636,798) | 3,097,552,996<br>(1,429,984,528) | 97.47 (98.91) | 98.59 (99.61) | 52.61(51.05) |
| D<sub>1</sub> | 11,686,644<br>(9,523,900) | 1,764,683,244<br>(1,419,937,618) | 96.71 (99.17) | 98.00 (99.69) | 46.47(45.7) |
| D<sub>2</sub> | 21,932,952<br>(16,115,664) | 3,311,875,752<br>(2,366,211,499) | 92.07 (98.26) | 95.01 (99.3) | 46.06(45.56) |
| D<sub>3</sub> | 16,077,412<br>(10,802,752) | 2,427,689,212<br>(1,606,861,271) | 96.88 (99.03) | 98.18 (99.64) | 48.66 (47.06) |
| **Total** | 105,222,908<br>(65,987,618) | 15,888,659,108<br>(9,752,530,317) | 95.44 (98.73) | 97.22 (99.52) | 49.17 (47.98) |

*Note: Values in each cell represent raw data point (data point after trimming and filtering).*

### Screening of differentially expressed genes (DEGs) in dwarf RILs

The FPKM values of each transcript for each of the 6 RIL libraries were calculated. Differential expression was observed in 22,897 genes (Fig 3A), although only 1,265 were found to be significantly differentially expressed (p<0.05) (Supplementary Fig 1 and Supplementary table T1). When the RILs were ranked from 1 to 6, N2 and N3 showed a somewhat similar number of upregulated (Rank 4 to 6) as well as downregulated (Rank 1 to 3) DEGs. A similar pattern was observed for the D2 and D3 RILs, although tall RILs had more upregulated DEGs (Fig 3B). All four RILs (N2, N3, D2, and D3) were the descendants of parents with their own phenotype; however, N1 and D1 were newly segregated from a parent of normal phenotype (Fig 1), which might explain the slightly different pattern of DEG ranks compared to their similar phenotype counterparts. Interestingly, N1 still had a slightly higher number of upregulated DEGs compared to downregulated ones while D1 DEGs showed the opposite pattern (Fig 3B).

**Fig 3.**
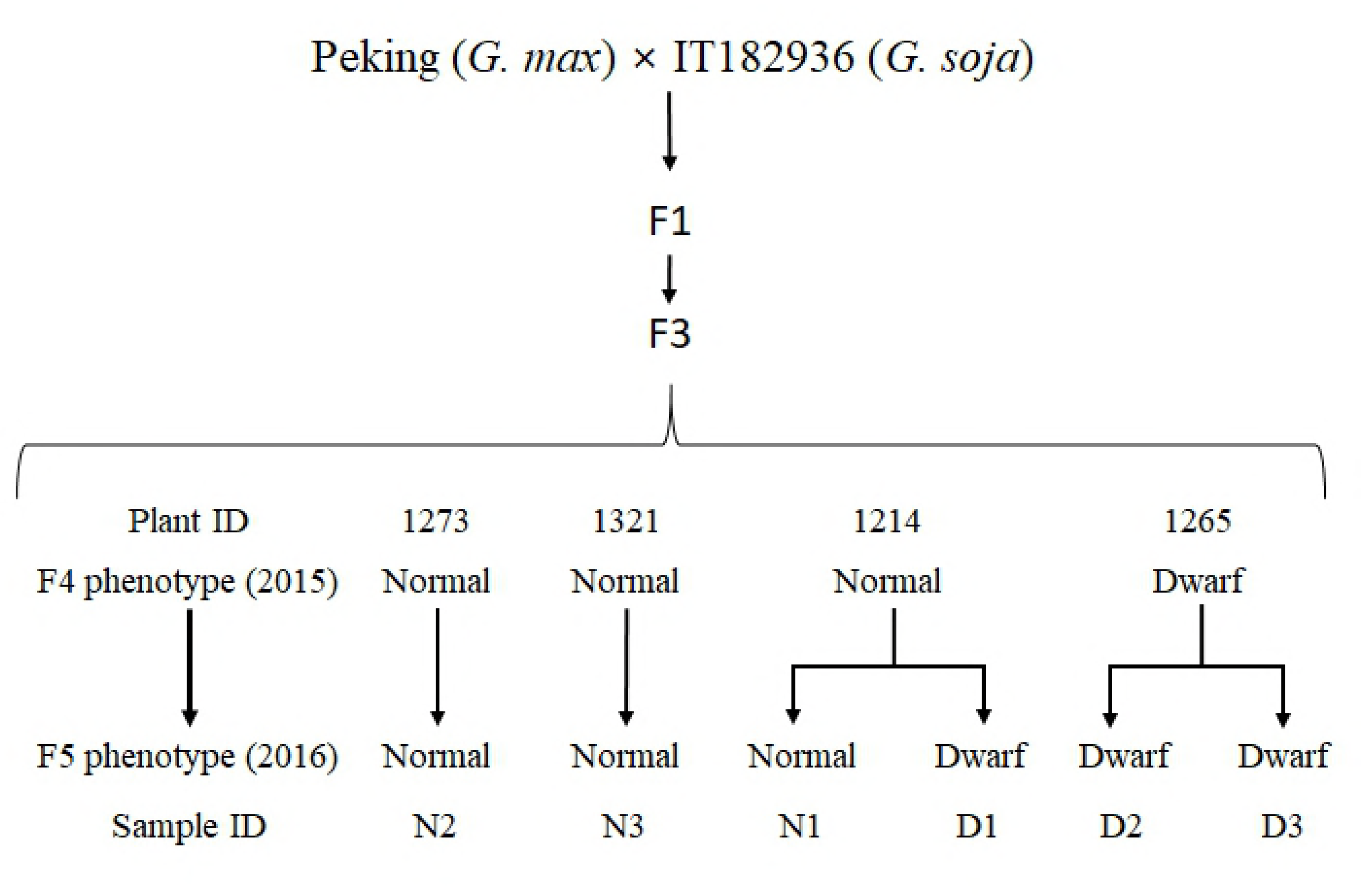
Preliminary analysis of differentially expressed genes (DEGs) in normal and dwarf RILs. A. Quantification of the DEGs according to their expression behavior. Green circle represents transcripts recorded in dwarf RILs while red one represents those recorded in normal RILs. Significantly up-regulated transcript numbers (p-value <0.05) are shown in non-overlapping section of each circles. B. DEGs annotated by GO terms of three different domains. BP= Biological Process; CC= Cellular component; MF= Molecular function C. Quantification of the DEGs in each of 6 RIL replicates according to their expression pattern. Rank-1 represents lowest expressing DEGs and Rank-6 encompasses highest expressing ones. The ranks were defined based on the level of expression of each DEG in the RILs.

### Categorization of DEGs according to GO terms

To gain further functional insights, the DEGs were categorically clustered under molecular function, biological process, and cellular component GOs, which assigned GO terms for only 912, 733, and 797 DEGs, respectively (Fig 3C). The total number of transcripts with at least one GO term was 1097. After removal of the genes with unassigned function, the DEGs were mainly assigned to ion binding (198 up- and 184 downregulated) under molecular function (Supplementary Table T1). However, when the differences in number of DEGs under the same GO term that were upregulated and downregulated were compared, dwarf RILs had the largest positive difference for the DEGs under kinase activity and largest negative difference for those under oxidoreductase. Other major GOs with larger numbers of upregulated DEGs included nucleic acid-binding transcription factor (TF), ion binding, signal transducer, and DNA binding. Similarly, other GOs with lower numbers of upregulated DEGs included molecular function, lyase activity, isomerase activity, transmembrane transporter, and peptidase activity. However, some GOs such as glycosyl bond hydrolase and methyl transferase had the same number of genes (12 and 10 respectively) in both up- and downregulated groups (Fig 3-A). Similar analysis of cellular component GOs showed the largest number of DEGs under the term ‘cellular component’ (251 up- and 270 downregulated) (Supplementary Table T1). A larger portion of the sequences were found to be downregulated in most of the GO terms. Major terms with lower numbers of upregulated DEGs included plastid, thylakoid, protein complex, cellular component, mitochondria, and vacuole Golgi apparatus. Other major terms with higher numbers of upregulated DEGs included plasma membrane, cytoplasm, extracellular region intracellular, and organelle. However, the differences of up- and downregulated genes in GO terms with higher numbers of downregulated DEGs were much wider than those in GO terms with higher numbers of upregulated DEGs (Fig 3-B). Additional clustering of the DEGs under biological process GO terms showed 54 different clusters among which all terms had higher numbers of downregulated DEGs compared to upregulated ones with the exception of only 12 GO terms. While most of the terms possessed few sequences, the GO term with highest DEGs was biosynthetic process (76 up- and 105 downregulated) (Supplementary Table T1). Interestingly, all of the DEGs under the photosynthesis GO term were downregulated. Other GO terms with comparatively higher numbers of downregulated DEGs included precursor and energy generation, small molecule metabolic process, biosynthetic process, cofactor metabolic process, biological process, and carbohydrate metabolic process. In contrast, The GO terms with relatively higher numbers of upregulated DEGs inlcuded response to stress, signal transduction, secondary metabolic process, and vesicle mediated transport. (Fig 3C).

### Pathway analysis of the DEGs

Among a total 1,097 DEGs with at least one GO term, KEGG pathway analysis revealed that only 281 were assigned to one or more enzyme IDs of 104 different pathways (Supplementary table T2). Among them, only the pathways with >5 DEGs were selected for further analysis in the present study. Up- or downregulated DEGs under each of these pathways were enumerated to see whether the pathways were positively or negatively affected in dwarf RILs (Fig 4). Even though the antibiotic biosynthesis pathway had the highest number of enzymes, the DEGs number was the highest in the purine metabolism pathway. None of the pathways had all of its genes upregulated; however, all DEGs in the pathways of ‘fructose metabolism’, ‘nitrogen metabolism’, ‘porphyrin and chlorophyll metabolism’, methane metabolism, and oxidative phosphorylation were downregulated. While most of the other pathways also had a comparatively larger proportion of downregulated DEGs (an additional 16 of the 33 pathways), pathways including ‘starch and sucrose metabolism’, ‘glutathione metabolism’, ‘phenylpropanoid biosynthesis’, ‘pyruvate metabolism’, ‘drug metabolism-cytochrome P450’, and thiamine metabolism had a relatively higher number of upregulated DEGs. Interestingly, ‘thiamine metabolism’ had the lowest enzyme-to-DEG ratio with 35 DEGs representing just a single enzyme (nucleoside-triphosphate phosphatase, EC 3.6.1.15).

**Fig 4.** Distribution of significantly up- or down-regulated genes clustered under KEGG pathways at level 3. Only the pathways with ≥5 enzymes or >6 HGEDs are shown in the figure. Full list of KEGG pathways is provided in the supplementary table S2.

In summary, the distribution of upregulated and downregulated genes is given in Table 2. It was observed that higher numbers of growth-related genes were downregulated whereas higher numbers of stress-responsive and defense-related genes were upregulated in dwarf soybean (Table 2). With close observation, dwarf RILs had upregulated PAMP (pathogen-associated molecular pattern) and DAMP (damage-associated molecular pattern) receptors such as TIR-NBS-LRR genes, cysteine-rich RLK, protein kinase superfamily, and lectin protein kinase. Expression of MAPK (mitogen-activated protein kinase), multiple WRKY genes, and PR-proteins (pathogenesis-related proteins), the downstream enzymes of stress response, were also upregulated in these plants. However, one group of receptors, LRR family proteins, was found to be downregulated (Table 3). Interestingly, heat shock protein 70, which, in combination with LRR family protein, activates MAPK (mitogen-activated protein kinase) precursors, was significantly upregulated in these lines. In addition to these, dwarf RILs also showed the upregulated expression of NB-ARC domain containing proteins, senescence-associated gene, SNF2, Ca2+ binding EF hands family protein, Ca2+ ATPase 2, early-response to dehydration protein, lycopene cyclase, and glucose-6-phosphate dehydrogenase, among others (Table 3). However, the dwarf RILs also had multiple downregulated genes which were related to photosynthesis including Calvin-Benson cycle, glycolysis, TCA cycle, and chlorophyll metabolism (Supplementary table T3).

**Table 2.** Number of significantly differentially expressed genes in normal and dwarf Soybean RILs.

| Gene category | Number of genes | Up regulated | Down regulated |
| --- | --- | --- | --- |
| Growth related | 71 | 31 | 40 |
| Abiotic Stress response | 65 | 44 | 20 |
| Defense responsive | 122 | 97 | 25 |
| Photosynthesis | 144 | 0 | 144 |
| Other categories | 863 | 344 | 519 |
| Total | 1265 |  |  |

**Table 3.** Significantly up and down regulated genes associated with various growth and defense genes. Up regulation is represented by positive Log2foldchange values whereas down regulation by negative Log2foldchange values.

| ID | log2FoldChange | Description |
| --- | --- | --- |
| Glyma.07G043000.2 | 3.4925 | Dormancy/auxin associated family protein |
| Glyma.08G339200.1 | 4.6926 | AP2/B3-like transcriptional factor family protein |
| Glyma.10G036700.1 | 3.1313 | ethylene response factor 1 |
| Glyma.12G103500.1 | 2.9116 | Auxin-responsive GH3 family protein |
| Glyma.15G070100.4 | 1.9970 | SNF2 domain-containing protein / helicase domain-containing protein / zinc finger protein-related |
| Glyma.15G091000.2 | 4.9711 | auxin response factor 8 |
| Glyma.15G091000.4 | 1.6981 | auxin response factor 8 |
| Glyma.16G050500.1 | 1.4788 | auxin signaling F-box 3 |
| Glyma.01G078200.1 | -3.7428 | SAUR-like auxin-responsive protein family |
| Glyma.06G158700.1 | -3.9887 | SAUR-like auxin-responsive protein family |
| Glyma.07G217900.1 | -4.3986 | Auxin efflux carrier family protein |
| Glyma.08G168300.1 | -3.2347 | SAUR-like auxin-responsive protein family |
| Glyma.11G126500.2 | -4.2138 | SKP1/ASK-interacting protein 16 |
| Glyma.12G011400.1 | -5.7415 | cytokinin oxidase/dehydrogenase 6 |
| Glyma.13G233900.1 | -4.1678 | ethylene responsive element binding factor 1 |
| Glyma.14G094800.1 | -2.6000 | cytokinin-responsive GATA factor 1 |
| Glyma.18G040000.1 | -2.2356 | RGA-like 2 |
| Glyma.18G091600.1 | -3.1957 | ethylene responsive element binding factor 3 |
| Glyma.01G032400.1 | 3.3612 | disease resistance protein (TIR-NBS-LRR class), putative |
| Glyma.01G128100.1 | 4.0166 | WRKY DNA-binding protein 33 |
| Glyma.03G087600.1 | 1.7825 | Disease resistance protein (TIR-NBS-LRR class) family |
| Glyma.04G218700.1 | 3.0687 | WRKY DNA-binding protein 51 |
| Glyma.06G123800.1 | 4.0283 | pathogenesis-related family protein |
| Glyma.06G258200.1 | 3.5113 | S-locus lectin protein kinase family protein |
| Glyma.06G263900.1 | 3.4751 | disease resistance protein (TIR-NBS-LRR class), putative |
| Glyma.06G265400.1 | 3.0768 | disease resistance protein (TIR-NBS-LRR class), putative |
| Glyma.07G007800.1 | 3.1485 | disease resistance protein (TIR-NBS-LRR class), putative |
| Glyma.07G037000.4 | 5.1035 | disease resistance protein (TIR-NBS-LRR class), putative |
| Glyma.07G057400.1 | 2.8571 | WRKY family transcription factor |
| Glyma.07G067900.1 | 2.0598 | disease resistance protein (TIR-NBS-LRR class), putative |
| Glyma.07G255400.2 | 3.3812 | mitogen-activated protein kinase 16 |
| Glyma.07G255400.3 | 3.3812 | mitogen-activated protein kinase 16 |
| Glyma.07G262700.1 | 2.4861 | WRKY family transcription factor |
| Glyma.08G352200.1 | 3.7378 | S-locus lectin protein kinase family protein |
| Glyma.10G138300.3 | 6.5765 | WRKY family transcription factor |
| Glyma.10G253100.1 | 2.5842 | cysteine-rich RLK (RECEPTOR-like protein kinase) 26 |
| Glyma.10G253200.1 | 1.7481 | cysteine-rich RLK (RECEPTOR-like protein kinase) 26 |
| Glyma.10G253300.1 | 2.3114 | cysteine-rich RLK (RECEPTOR-like protein kinase) 29 |
| Glyma.10G253300.2 | 2.3114 | cysteine-rich RLK (RECEPTOR-like protein kinase) 29 |
| Glyma.10G253900.1 | 2.3413 | cysteine-rich RLK (RECEPTOR-like protein kinase) 26 |
| Glyma.11G152200.1 | 2.0371 | S-locus lectin protein kinase family protein |
| Glyma.11G207000.2 | 2.4919 | cysteine-rich RLK (RECEPTOR-like protein kinase) 2 |
| Glyma.11G207500.1 | 2.1918 | cysteine-rich RLK (RECEPTOR-like protein kinase) 2 |
| Glyma.12G073000.1 | 2.0150 | mitogen-activated protein kinase 3 |
| Glyma.12G132000.1 | 2.7420 | disease resistance protein (TIR-NBS-LRR class), putative |
| Glyma.12G135600.3 | 4.7007 | disease resistance protein (TIR-NBS-LRR class), putative |
| Glyma.12G212500.1 | 3.3779 | disease resistance protein (TIR-NBS-LRR class), putative |
| Glyma.13G117600.1 | 2.7217 | WRKY family transcription factor |
| Glyma.13G303800.1 | 2.0579 | S-locus lectin protein kinase family protein |
| Glyma.13G370100.1 | 2.8501 | WRKY DNA-binding protein 40 |
| Glyma.14G102900.1 | 3.4983 | WRKY DNA-binding protein 40 |
| Glyma.14G103100.1 | 3.2818 | WRKY DNA-binding protein 40 |
| Glyma.15G062400.1 | 3.1915 | basic pathogenesis-related protein 1 |
| Glyma.15G062500.1 | 2.6191 | basic pathogenesis-related protein 1 |
| Glyma.15G062700.1 | 7.2652 | basic pathogenesis-related protein 1 |
| Glyma.15G186300.1 | 3.6211 | WRKY family transcription factor |
| Glyma.15G213600.1 | 3.0120 | S-locus lectin protein kinase family protein |
| Glyma.16G026400.1 | 2.2147 | WRKY family transcription factor |
| Glyma.16G085700.1 | 2.0813 | Disease resistance protein (TIR-NBS-LRR class) family |
| Glyma.16G159300.1 | 3.4444 | Disease resistance protein (TIR-NBS-LRR class) family |
| Glyma.16G213700.1 | 5.1618 | Disease resistance protein (TIR-NBS-LRR class) family |
| Glyma.16G213900.1 | 4.5921 | Disease resistance protein (TIR-NBS-LRR class) family |
| Glyma.17G222500.1 | 3.8782 | WRKY DNA-binding protein 40 |
| Glyma.17G224200.1 | 3.7199 | Concanavalin A-like lectin protein kinase family protein |
| Glyma.17G224300.2 | 2.7373 | Concanavalin A-like lectin protein kinase family protein |
| Glyma.18G046300.1 | 2.9368 | cysteine-rich RLK (RECEPTOR-like protein kinase) 2 |
| Glyma.18G056600.1 | 2.6815 | WRKY DNA-binding protein 33 |
| Glyma.18G081200.4 | 4.0941 | WRKY family transcription factor family protein |
| Glyma.18G219700.1 | 4.7267 | cysteine-rich RLK (RECEPTOR-like protein kinase) 23 |
| Glyma.20G020200.1 | 2.5115 | disease resistance protein (TIR-NBS-LRR class), putative |
| Glyma.20G061300.1 | 3.9699 | disease resistance protein (TIR-NBS-LRR class), putative |
| Glyma.20G137700.1 | 2.7652 | cysteine-rich RLK (RECEPTOR-like protein kinase) 29 |
| Glyma.20G137900.1 | 3.8608 | cysteine-rich RLK (RECEPTOR-like protein kinase) 29 |
| Glyma.20G138500.1 | 2.6261 | cysteine-rich RLK (RECEPTOR-like protein kinase) 29 |
| Glyma.20G140200.1 | 2.3726 | cysteine-rich RLK (RECEPTOR-like protein kinase) 25 |
| Glyma.20G140400.1 | 2.3326 | cysteine-rich RLK (RECEPTOR-like protein kinase) 25 |
| Glyma.20G140500.1 | 3.2787 | cysteine-rich RLK (RECEPTOR-like protein kinase) 10 |
| Glyma.14G031300.1 | 5.3553 | NPR1-like protein 3 |
| Glyma.01G244600.1 | -4.5223 | Leucine-rich repeat (LRR) family protein |
| Glyma.08G152800.1 | -3.8174 | Leucine-rich repeat (LRR) family protein |
| Glyma.08G152800.2 | -3.3731 | Leucine-rich repeat (LRR) family protein |
| Glyma.13G269100.1 | -3.8839 | pathogenesis-related family protein |
| Glyma.13G341500.1 | -5.0925 | Leucine-rich repeat (LRR) family protein |

In addition to the upregulation of stress-related DEGs and downregulation of primary metabolism-related genes, some transcripts involved in hormone signaling also showed significantly differential expression in dwarf RILs. The plants showed the downregulation of SAUR-like auxin-responsive proteins and upregulation of auxin response factor 8, auxin-responsive GH3 family protein, auxin signaling F-box, and auxin-associated family protein. Others, such as RGL2 (RGA-like 2, a GA signaling repressor), SKP1, cytokinin-responsive GATA factor, cytokinin oxidase, and ethylene responsive element binding factor were downregulated while DEGs encoding ethylene response factor, AP2/B3-like transcriptional factor, ERD (early-responsive to dehydration) protein, and SNF2 domain containing protein, among others, were significantly upregulated (Fig 5). Moreover,the majority of the uniquely significant DEGs (with all of their versions expressing significantly) were photosynthesis-related (upregulated in normal RILs) and disease/stress-related (upregulated in dwarf RILs) (Fig 5).

**Fig 5.** Selected heatmaps of DEGs. A) Expression of the DEGs involved in PAMP receptors. B) Expression pattern of the DEGs active in salicylic acid (SA) signaling. C) Expression pattern of genes related to plant immunity response. D) Expression pattern of genes involved in chloroplast metabolism and thylakoid reaction, one of the reactions of photosynthesis. E) Expression pattern of glycolysis and pentose phosphate pathway associated genes

### Salicylic acid determination by HPLC

Observing the upregulated state of stress/disease-responsive DEGs in dwarf RILs (Fig 7), we chose to assess the change in plant salicylic acid (SA) content. HPLC analysis was conducted for different tall and dwarf RILs along with their inital parents (*G. max* cv. Peking and *G. soja*). The result clearly showed higher SA contents in dwarf RILs compared to their tall counterparts (Fig 6). Additionally, *G. soja* showed relatively higher SA content than *G. max* suggesting a relatively highly activated state of SA signaling in the latter. Due to the short stature of the dwarf RILs and putatively activated SA signaling in the plants, the plants were tested for possible infection by soybean dwarf virus. The analysis gave negative results suggesting that the short stature of dwarf RILs was clearly due to genetic segregation and not because of viral infection.

**Fig 6.** HPLC analysis of *G. max* vs Peking, *G. soja* and their normal (N1-N3) and dwarf (D1-D3) F5 RIL progenies for their salicylic acid (SA) content.

## Discussion

### Dwarf RILs have upregulated stress responsive genes

Stresses are known to counteract overall plant growth. In the present study, we found the largest number of disease-responsive genes and stress-responsive genes upregulated in dwarf soybean RILs, suggesting their constitutive expression. However, the KEGG pathway showed that the number of upregulated DEGs involved in antibiotic biosynthesis were fewer than the downregulated ones, indicating comparatively lower immunity of the dwarf RILs. Overexpression of disease resistance and other immune responsive genes has been attributed to produce an overdose effect in plants, thereby activating the downstream defense response pathways and curtailing plant growth by inhibiting protein synthesis [26, 27]. Huot et al. [3] also strongly argued for the reduced growth of plants under pathogenic pressure. The auto-toxicity of constitutively expressed defense-responsive gene products hindering plant growth has also been reported in tobacco [28]. In a separate extensive study on wheat, Heil et al. [29] strongly argued for the tradeoff between the defense response and plant growth because of the energetically costly molecular mechanism. Programmed cell death (PCD) induced by the constitutive expression of pathogenesis-related genes is a general phenomenon in plants [30]. The present study also showed the significant upregulation of a DCD (development and cell death) domain protein in dwarf RILs, likely in response to the upregulated disease responsive genes. The gene, as its name suggests, is involved in development and PCD in plants [31]. Severe dwarfism was also reported in *Arabidopsis thaliana* due to the disruption of a MAP3K gene (MEKK1), which led to constitutive expression of pathogenesis-associated genes in the plant [32].

### Genes involved in hormonal regulation

Auxin plays a vital role in plant growth and development [33]. Our dwarf RILs showed upregulated auxin response factor (ARF8) and auxin-responsive GH3 family protein. Interestingly, the GH3 gene has been reported to be involved in negative regulation of shoot cell elongation and lateral root formation [34]. Moreover, ARF8 is known to play a role in controlling auxin (IAA) levels through negative feedback by regulating GH3, thereby reducing hypocotyl growth [35]. Additionally, the small auxin-up RNA (SAUR)-like gene was found to be downregulated in the dwarf RILs. As the gene is reportedly regulated by the auxin level in plants and is involved in cell division [36], it is likely that the dwarf RILs have comparatively lower auxin content than their tall counterparts. Another study by ten Have [37] showed resistance to higher IAA concentrations by LRR RLK mutant *Arabidopsis thaliana* plants and suggested the involvement of LRR RLK genes in auxin signaling. Interestingly, the dwarf soybean RILs in the present study showed down regulation of the LRR family protein (Fig 5) which might have hindered auxin signaling in the plants.

### Dwarf phenotype is likely due to a hypersensitive response

Salicylic acid (SA) is a signaling molecule and is generally associated with biotic stress in plants [38, 39]. SA is necessary for the plant’s systemic acquired resistance (SAR) [40]. The molecule brings about a hypersensitive response (HR) when accumulated in plants with high reaction oxygen species and actively expressed pathogenesis-related (PR) genes [41]. Our study showed that dwarf soybean RILs had higher SA content (Fig 6) in addition to the upregulated states of different pathogen-associated molecular pattern (PAMP) receptor genes (Fig 5A and 5B). As SA signaling in plants under stress precedes ROS bursts in different intracellular compartments [42], it is highly likely that the dwarf RILs also have high ROS. This phenomenon is supported, in part, by the upregulated state of respiratory burst oxidase in the dwarf RILs (Supplementary table T1). The gene is also reported to **be involved in disease resistance** [43]. Additionally, the comparatively higher SA in *G. max* relative to that in *G. soja* suggests the higher expression of PR genes in the former. In general, HR is characterized by the necrosis at the site of pathogenesis [40]. Since there was no apparent pathogenesis in the plants (dwarf RILs), the accumulated SA might instead have contributed to stunting of the plants.

### Genes involved in energy metabolism are downregulated in dwarf RILs

Our study showed that dwarf RILs had relatively lower expression of both nitrate reductase (NR) and nitrite reductase (NiR). These genes are closely associated with nitrate assimilation as NR reduces nitrate to nitrite that, in turn, is reduced to ammonia by NiR, and ammonia is finally fixed into carbon [44]. Additionally, the study by **Crawford** has reported the severely dwarf phenotype of NiR-antisense tobacco plants [44].

In normal healthy plants, glucose is metabolized via glycolysis and the non-oxidative branch of the pentose phosphate pathway (PPP) [45-47]. However, in plants under stress, the oxidative branch of PPP is more active which is critical to drive the conversion of glucose 6-phosphate to CO2, ribulose-5- phosphate, and NADPH [47]. The present study found the clear upregulation of glucose-6-phosphate dehydrogenase, a key enzyme that mediates the reaction, indicating the possible constitutive stress/defense response in the plants at the molecular level. Moreover, the key enzymes of the Calvin-Benson cycle and chlorophyll metabolism were found to be exclusively downregulated, suggesting a lower rate of primary metabolism in the dwarf RILs. Such global downregulation of genes involved in photosynthesis is common in plants under biotic stress [38]. However, their lower states of expression in dwarf RILs compared to their tall counterparts (even though all plants were grown under the same conditions) also indicate a constitutively active state of biotic defense genes in the dwarf RILs. An earlier study [48] also reported the decreased expression of genes involved in photosynthesis, in contrast to increased expression of defense-related genes, upon herbivory in tobacco. Moreover, overall downregulation of photosynthesis-related genes is common in plants which are under biotic stress as the plants have to re-allocate their resources from growth to defense [38].

### Conclusion

We obtained the segregation of RILs in dwarf and normal lines. There were highly significantly up/downregulated genes in the comparison of gene expression in normal and dwarf soybeans. We observed that dwarf lines exhibited higher expression of stress- and defense-related genes. We also found higher levels of SA in dwarf lines, which may have contributed to their stunted growth phenotype. Such over-expression of disease resistance and other immune responsive genes were targeted to understand how the immune genes regulate the response of plant growth. In addition, all the photosynthesis-related genes were downregulated in dwarf plants. The transcriptome expression and genes classified as related to plant growth may be useful resources for researchers studying plant growth.

### Author Contribution

I.-Y.C conceived and designed the project and prepared the samples. Y.-W.B and N.S.R were major contibutor to the data anaysis and writing the manuscript. H.Y analyzed the metabolite. H.-K.C and J.-H.K, P.B and K.C.P surveyed and analyzed the NGS data. K.-S. H and J.-K. H cultivated and surveyed the soybean population.

## Acknowledgement

This work was supported by the National Research Foundation of Korea (NRF) grant funded by the Korea government (MSIT) (No. 2017R1A2B4011198) and by a 2017 Research Grant from Kangwon National University.

## References

1. Acquaah G. Principles of plant genetics and breeding: John Wiley & Sons; 2009.

2. Peng J, Richards DE, Hartley NM, Murphy GP, Devos KM, Flintham JE, et al. 'Green revolution' genes encode mutant gibberellin response modulators. Nature. 1999; 400: 256-61.

3. Huot B, Yao J, Montgomery BL, He SY. Growth-defense tradeoffs in plants: a balancing act to optimize fitness. Mol Plant. 2014; 7: 1267-87.

4. Hutchings MJ, de Kroon H. Foraging in Plants: the Role of Morphological Plasticity in Resource Acquisition. Advances in ecological research. Advances in Ecological Research. 25: Elsevier; 1994. p. 159-238.

5. Boutraa T, Akhkha A, Al-Shoaibi AA, Alhejeli AM. Effect of water stress on growth and water use efficiency (WUE) of some wheat cultivars (Triticum durum) grown in Saudi Arabia. J Taibah Univ Sci. 2010; 3: 39-48.

6. Tatrai ZA, Sanoubar R, Pluhar Z, Mancarella S, Orsini F, Gianquinto G. Morphological and Physiological Plant Responses to Drought Stress in *Thymus citriodorus*. International Journal of Agronomy. 2016; 2016:

7. Wang X, Deng Z, Zhang W, Meng Z, Chang X, Lv M. Effect of Waterlogging Duration at Different Growth Stages on the Growth, Yield and Quality of Cotton. PLoS One. 2017; 12: e0169029.

8. Striker GG. Flooding stress on plants : anatomical, morphological and physiological responses. Botany. 2012; 3-28.

9. Korner C. Plant adaptation to cold climates. F1000Research. 2016; 5:

10. Daft MJ, Nicolson T. Effect of Endogone mycorrhiza on plant growth. New Phytologist. 1966; 65: 343-50.

11. Scholthof HB, Scholthof KB, Jackson AO. Identification of tomato bushy stunt virus host-specific symptom determinants by expression of individual genes from a potato virus X vector. Plant Cell. 1995; 7: 1157-72.

12. Poveda K, Steffan-Dewenter I, Scheu S, Tscharntke T. Effects of below- and above-ground herbivores on plant growth, flower visitation and seed set. Oecologia. 2003; 135: 601-5.

13. Cooper R, Martin R, Schmitthenner A, McBlain B, Fioritto R, St Martin S, et al. Registration of'Hobbit 87'soybean. Crop science (USA). 1991; 31: 1093.

14. Cooper R, Martin R, St Martin S, Calip-DuBois A, Fioritto R, Schmitthenner A. Registration of' Charleston'soybean. Crop science. 1995; 35: 593.

15. Cooper R, Mendiola T, Martin SS, Fioritto R, Schmitthenner A, Dorrance A. Registration ofStrong'Soybean. Crop science. 2001; 41: 921.

16. Cooper R, Mendiola T, St Martin S, Fioritto R, Dorrance A. Registration of' Apex'soybean.(Registration Of Cultivars). Crop Science. 2003; 43: 1563-4.

17. de Souza IR, MacAdam JW. Gibberellic acid and dwarfism effects on the growth dynamics of B73 maize (Zea mays L.) leaf blades: a transient increase in apoplastic peroxidase activity precedes cessation of cell elongation. J Exp Bot. 2001; 52: 1673-82.

18. Zhang Y, Turner JG. Wound-induced endogenous jasmonates stunt plant growth by inhibiting mitosis. PLoS One. 2008; 3: e3699.

19. Ockerse R, Galston AW. Gibberellin-auxin interaction in pea stem elongation. Plant Physiol. 1967; 42: 47-54.

20. Tanimoto E. Tall or short? Slender or thick? A plant strategy for regulating elongation growth of roots by low concentrations of gibberellin. Ann Bot. 2012; 110: 373-81.

21. Zhang P, Liu X, Tong H, Lu Y, Li J. Association mapping for important agronomic traits in core collection of rice (Oryza sativa L.) with SSR markers. PLoS One. 2014; 9: e111508.

22. Hwang WJ, Kim MY, Kang YJ, Shim S, Stacey MG, Stacey G, et al. Genome-wide analysis of mutations in a dwarf soybean mutant induced by fast neutron bombardment. Euphytica. 2015; 203: 399-408.

23. Bolger AM, Lohse M, Usadel B. Trimmomatic: a flexible trimmer for Illumina sequence data. Bioinformatics. 2014; 30: 2114-20.

24. Kim D, Langmead B, Salzberg SL. HISAT: a fast spliced aligner with low memory requirements. Nat Methods. 2015; 12: 357-60.

25. Li H, Durbin R. Fast and accurate short read alignment with Burrows-Wheeler transform. Bioinformatics. 2009; 25: 1754-60.

26. Belkhadir Y, Subramaniam R, Dangl JL. Plant disease resistance protein signaling: NBS–LRR proteins and their partners. Current opinion in plant biology. 2004; 7: 391-9.

27. Tao Y, Yuan F, Leister RT, Ausubel FM, Katagiri F. Mutational analysis of the Arabidopsis nucleotide binding site-leucine-rich repeat resistance gene RPS2. Plant Cell. 2000; 12: 2541-54.

28. Baldwin IT, Callahan P. Autotoxicity and chemical defense: nicotine accumulation and carbon gain in solanaceous plants. Oecologia. 1993; 94: 534-41.

29. Heil M, Hilpert A, Kaiser W, Linsenmair KE. Reduced growth and seed set following chemical induction of pathogen defence: does systemic acquired resistance (SAR) incur allocation costs? Journal of Ecology. 2000; 88: 645-54.

30. Greenberg JT. Programmed Cell Death in Plant-Pathogen Interactions. Annu Rev Plant Physiol Plant Mol Biol. 1997; 48: 525-45.

31. Tenhaken R, Doerks T, Bork P. DCD - a novel plant specific domain in proteins involved in development and programmed cell death. BMC Bioinformatics. 2005; 6: 169.

32. Suarez-Rodriguez MC, Adams-Phillips L, Liu YD, Wang HC, Su SH, Jester PJ, et al. MEKK1 is required for flg22-induced MPK4 activation in Arabidopsis plants. Plant Physiol. 2007; 143: 661-9.

33. Quint M, Gray WM. Auxin signaling. Curr Opin Plant Biol. 2006; 9: 448-53.

34. Nakazawa M, Yabe N, Ichikawa T, Yamamoto YY, Yoshizumi T, Hasunuma K, et al. DFL1, an auxin-responsive GH3 gene homologue, negatively regulates shoot cell elongation and lateral root formation, and positively regulates the light response of hypocotyl length. Plant J. 2001; 25: 213-21.

35. Tian CE, Muto H, Higuchi K, Matamura T, Tatematsu K, Koshiba T, et al. Disruption and overexpression of auxin response factor 8 gene of Arabidopsis affect hypocotyl elongation and root growth habit, indicating its possible involvement in auxin homeostasis in light condition. Plant J. 2004; 40: 333-43.

36. Markakis MN, Boron AK, Van Loock B, Saini K, Cirera S, Verbelen JP, et al. Characterization of a small auxin-up RNA (SAUR)-like gene involved in Arabidopsis thaliana development. PLoS One. 2013; 8: e82596.

37. ten Hove CA, Bochdanovits Z, Jansweijer VM, Koning FG, Berke L, Sanchez-Perez GF, et al. Probing the roles of LRR RLK genes in Arabidopsis thaliana roots using a custom T-DNA insertion set. Plant Mol Biol. 2011; 76: 69-83.

38. Bilgin DD, Zavala JA, Zhu J, Clough SJ, Ort DR, DeLucia EH. Biotic stress globally downregulates photosynthesis genes. Plant Cell Environ. 2010; 33: 1597-613.

39. Herrera-Vasquez A, Salinas P, Holuigue L. Salicylic acid and reactive oxygen species interplay in the transcriptional control of defense genes expression. Front Plant Sci. 2015; 6: 171.

40. Dempsey DMA, Shah J, Klessig DF. Salicylic Acid and Disease Resistance in Plants. Critical Reviews in Plant Sciences. 1999; 18: 547-75.

41. Lam E, Kato N, Lawton M. Programmed cell death, mitochondria and the plant hypersensitive response. Nature. 2001; 411: 848-53.

42. Wrzaczek M, Brosche M, Kangasjarvi J. ROS signaling loops - production, perception, regulation. Curr Opin Plant Biol. 2013; 16: 575-82.

43. Pratap A, Kumar J. Alien Gene Transfer in Crop Plants, Volume 22014.

44. Crawford NM. Nitrate: nutrient and signal for plant growth. Plant Cell. 1995; 7: 859-68.

45. Bogorad IW, Lin TS, Liao JC. Synthetic non-oxidative glycolysis enables complete carbon conservation. Nature. 2013; 502: 693-7.

46. Morot-Gaudry-Talarmain Y, Rockel P, Moureaux T, Quillere I, Leydecker MT, Kaiser WM, et al. Nitrite accumulation and nitric oxide emission in relation to cellular signaling in nitrite reductase antisense tobacco. Planta. 2002; 215: 708-15.

47. Stincone A, Prigione A, Cramer T, Wamelink MM, Campbell K, Cheung E, et al. The return of metabolism: biochemistry and physiology of the pentose phosphate pathway. Biol Rev Camb Philos Soc. 2015; 90: 927-63.

48. Giri AP, Wunsche H, Mitra S, Zavala JA, Muck A, Svatos A, et al. Molecular interactions between the specialist herbivore Manduca sexta (Lepidoptera, Sphingidae) and its natural host Nicotiana attenuata. VII. Changes in the plant's proteome. Plant Physiol. 2006; 142: 1621-41.

